# A quantitative framework for motion visibility in human cortex

**DOI:** 10.1101/359562

**Authors:** Daniel Birman, Justin L Gardner

**Affiliations:** Department of Psychology, Stanford University, Stanford, CA 94305

## Abstract

Despite the central use of motion visibility to reveal the neural basis of perception, perceptual decision making, and sensory inference there exists no comprehensive quantitative framework establishing how motion visibility parameters modulate human cortical response. Random-dot motion stimuli can be made less visible by reducing image contrast or motion coherence, or by shortening the stimulus duration. Because each of these manipulations modulates the strength of sensory neural responses they have all been extensively used to reveal cognitive and other non-sensory phenomenon such as the influence of priors, attention, and choice-history biases. However, each of these manipulations is thought to influence response in different ways across different cortical regions and a comprehensive study is required to interpret this literature. Here, human participants observed random-dot stimuli varying across a large range of contrast, coherence, and stimulus durations as we measured blood-oxygen-level dependent responses. We developed a framework for modeling these responses which quantifies their functional form and sensitivity across areas. Our framework demonstrates the sensitivity of all visual areas to each parameter, with early visual areas V1-V4 showing more parametric sensitivity to changes in contrast and V3A and MT to coherence. Our results suggest that while motion contrast, coherence, and duration share cortical representation, they are encoded with distinct functional forms and sensitivity. Thus, our quantitative framework serves as a reference for interpretation of the vast perceptual literature manipulating these parameters and shows that different manipulations of visibility will have different effects across human visual cortex and need to be interpreted accordingly.

## New & Noteworthy

Manipulations of motion visibility have served as a key tool for understanding the neural basis for visual perception, decision-making, and sensory inference. However, knowledge about motion visibility representation in human cortex is sparse. Here we measured human cortical response to changes in visibility across a comprehensive range of motion visibility parameters (contrast, coherence, and duration) and modeled these with a quantitative framework. We find that all retinotopically-defined visual areas were sensitive to these parameters, but differed significantly in the magnitude of sensitivity and form of relationship. Our quantitative framework can be used as a reference for linking human cortical response to perception and underscores that different manipulations of motion visibility can have greatly different effects on cortical representation.

## Introduction

Much of the neural basis of perception has been revealed by manipulations that control the visibility of motion stimuli. For example, global motion direction of random-dot stimuli is made less visible by decreasing motion coherence, i.e. the percentage of dots moving in the same direction. At lower visibility levels, small changes in cortical signals manifest in measurable behavioral effects, thus documenting direct links between cortical physiology and perception (Newsome et al. 1989; Britten et al. 1992) and uncovering neural signals supporting evidence accumulation (Shadlen et al. 1996; Shadlen and Newsome 2001; Roitman and Shadlen 2002; Huk and Shadlen 2005; Katz et al. 2016). Making stimuli brief also renders them less visible, aiding, for example, the study of information integration across eye movements (Melcher and Morrone 2003). Increasing image contrast, the average difference between bright and dark (Bex and Makous 2002), makes stimuli more visible and cortical responses monotonically larger allowing links to be made between cortical response and perception (Boynton et al. 1999; Ress et al. 2000; Ress and Heeger 2003), disambiguating mechanisms for spatial attention (Carrasco et al. 2000; Pestilli et al. 2011; Hara and Gardner 2014; Hara et al. 2014), uncovering neural correlates of conscious perception (Lumer et al. 1998; Wunderlich et al. 2005), and revealing the effects of putative priors (Stocker and Simoncelli 2006; Vintch and Gardner 2014). While each of these manipulations has been used extensively in the human perceptual literature, they can have greatly different effects on human neural response. Given the central importance of motion visibility a quantitative model of response across human visual cortex is required to provide a framework for interpreting and building upon these various findings.

Such a population response model must quantitatively account for the shape of the relationship between motion visibility and cortical response. The response function for contrast has been characterized as a sigmoidal function for measurements in single-unit (Albrecht and Hamilton 1982; Sclar et al. 1990) and populations (Tootell and Taylor 1995; Boynton et al. 1996, 1999; Tootell et al. 1998; Logothetis et al. 2001; Avidan et al. 2002; Olman et al. 2004; Gardner et al. 2005). Increasing motion coherence typically results in linear increases in response (Britten et al. 1993; Simoncelli and Heeger 1998; Rees et al. 2000; Aspell et al. 2005; Händel et al. 2007) although this may depend on the exact stimulus parameters (Ajina et al. 2015).

A population response model must also quantify the variable sensitivity to visibility parameters across cortical areas. The earliest cortical areas have a larger dynamic range for contrast compared to later areas which are more invariant (Rolls and Baylis 1986; Sclar et al. 1990; Cheng et al. 1994; Avidan et al. 2002). Less is known about motion coherence sensitivity except that the neural response to coherent compared to incoherent motion or blank evokes a large response in human MT with some sensitivity reported in earlier visual cortical areas (Zeki et al. 1991; Watson et al. 1993; Dupont et al. 1994; Tootell et al. 1995; Heeger et al. 1999; Costagli et al. 2014; Ajina et al. 2015) and parietal and ventral regions (Braddick et al. 2001).

Finally this model must account for stimulus duration effects. Hemodynamic responses to visual stimuli are approximately temporally linear except when durations (Boynton et al. 1996, 2012) or inter-stimulus intervals (Huettel and McCarthy 2000) are brief. The divergence from linearity may differ across cortical areas (Birn et al. 2001) and motion-sensitive regions may be most sensitive to transient changes (Stigliani et al. 2017). Here we measured blood-oxygen-level dependent (bold>) (Ogawa et al. 1990) response in human observers to a large range of contrast, coherence and duration of motion stimuli, and built a quantitative model linking these visibility properties with physiological response in retinotopically defined visual areas. Sensitivity to these parameters varied significantly across areas, although all were sensitivity to both contrast and coherence without interaction. While perceptual experiments have often used different means of affecting visibility interchangeably our results provide a reference model that underscores the differences in response to each manipulation of visibility across cortical areas, thus providing a quantifiable way to interpret experiments that link cortical response to perception.

## Methods

### Observers

In total, 11 observers (8 female, 3 male; mean age 26 y; age range 19-36 y) were subjects for the experiments. All observers except one (who was an author) were naïve to the intent of the experiments. Observers were scanned three times, in 2 two-hour sessions of the experiment and a one hour retinotopy session. Procedures were approved in advance by the Stanford Institutional Review Board on human participants research and all observers gave prior written informed consent before they participated in the experiment. When necessary, observers wore corrective lenses to correct their vision to normal.

### Hardware setup for stimulus and task control

Visual stimuli were generated using MATLAB (The Mathworks, Inc.) and MGL (Gardner et al. 2018b). Stimuli were back-projected via an Eiki LC-WUL100L projector (resolution of 1900×1200, refresh-rate of 100Hz) onto an acrylic sheet mounted inside the scanner bore near the head coil. Visual stimuli were viewed through a mirror mounted on the head coil and responses were collected via an MRI-compatible button box. Output luminance was measured with a PR650 spectrometer (Photo Research, Inc.) and a neutral density filter used to set the average screen luminance to 300 cd/m2. The gamma table was then dynamically adjusted at the beginning of each trial to linearize the luminance display such that the full 10-bit output resolution of the gamma table could be used to display the maximum contrast needed. Other sources of light were minimized during scanning.

### Eye tracking

Prior to the experiment subjects were extensively trained on a behavioral task requiring precise fixation. Eye-tracking was performed using an infrared video-based eye-tracker at 500 Hz (Eyelink 1000; SR Research). Calibration was performed throughout each session to maintain a validation accuracy of less than 1 degree average offset from expected using either a ten-point or thirteen-point calibration procedure. Trials were canceled on-line when observer’s eyes moved more than 1 degree away from the fixation cross for more than 300 ms. After training, canceled trials consisted of fewer than 0.1% of all trials. Due to technical limitations eye tracking was not performed inside the scanner.

### Experimental design

Motion stimuli consisted of two patches of moving dots and a central cross (1 x 1 deg) on which observers maintained fixation. The dot patches were rectangular regions extending from 3.5 to 12 deg horizontal and −7 to 7 deg vertical. Each patch was filled with 21 dots / deg^2^, 50% brighter and 50% darker than the gray background (300 cd/m^2). Both patches maintained a constant baseline in between trials of 25% contrast and incoherent motion. During a trial, the patches increased in either or both contrast and coherence. To minimize involuntary eye movements, the coherent dot motion direction was randomized to be horizontally inward or outward from fixation on each trial, such that each patch moved in opposite direction. All dots moved at 6 deg / s updated on each video frame. Motion strength was adjusted by changing motion coherence, that is, the percentage of dots that moved in a common direction with all other dots moving in random directions. Dots were randomly assigned on each video frame to be moving in the coherent or random directions.

We measured the cortical response to a wide range of brief increments of stimulus contrasts and coherences of variable durations while observers performed an independent and asynchronous task at fixation (Fig 1). Each scan began with a 30 s baseline period (25% contrast, 0% coherence) to allow visual cortex to adapt. Each trial consisted of a brief increment in either or both the contrast and motion coherence of the dot patches. The dot patches then returned to baseline (25% contrast, 0% coherence) for an inter-trial interval of 2 to 11 s (mean 6.5 s) randomly sampled from an exponential distribution. The next trial then began synchronized to the next volume acquisition of the magnet. Stimulus increments were chosen to be +0, +25, +50, or +75% above the baseline 25% contrast and +0, +25, +50, +75, or +100% above the baseline 0% coherence and lasted for 250, 500, 1000, 2000, 2500 or 4000 ms (or as close to these durations as the display frame refresh would allow). We presented trials in two sets; a “complete cross set” in which all combinations of contrast and coherence changes at 2500 ms duration were presented (4 contrasts x 5 coherences = 20 conditions) and a “duration set” in which a subset of the contrast and coherence combinations (+25 or +75 contrast and +25 or +100 coherence) were presented for variable stimulus durations (4 contrast and coherence combinations x 5 stimulus durations = 20 conditions). Thus, across the complete cross and duration sets, there was a total of 40 conditions (20 each in the complete cross and duration sets). For each condition we acquired a minimum of 20 repeated presentations throughout the scan sessions of each observer, resulting in a minimum of 800 trials total. The two trial sets were presented in separate scans interleaved within sessions. Condition order within each scan, for both trial sets, was randomized independently for the stimulus on the left and right such that in every block of 40 trials all conditions were presented in both dot patches.

**Figure 1.**
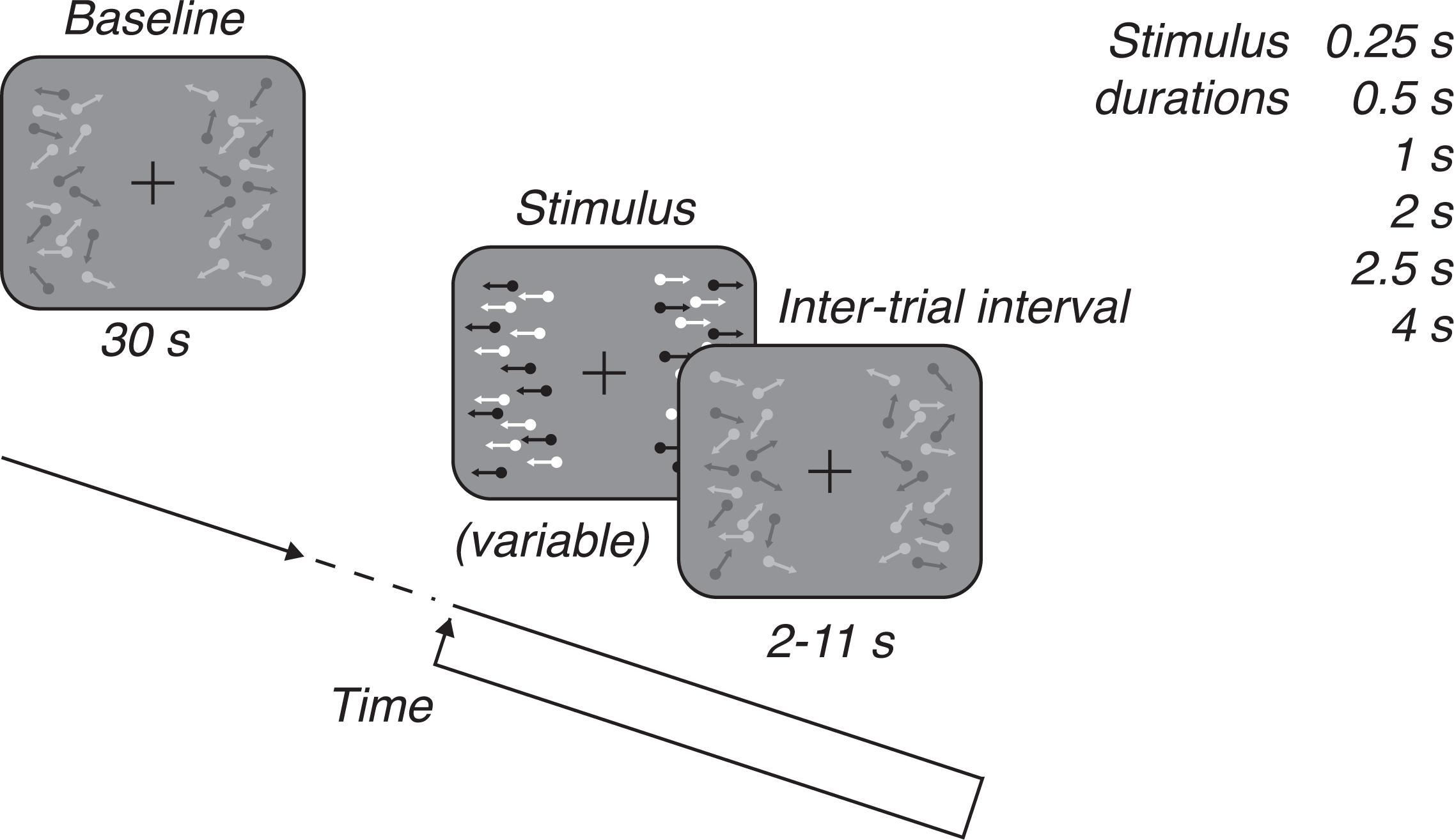
Cortical measurement experiment. Observers were shown patches of moving dots that increased in contrast and motion coherence on each trial. A 30 s baseline period preceded each scan with 25% contrast dots and incoherent motion and the baseline dots persisted between trials. On each trial the contrast increased by 0, 25, 50, or 75% and the coherence by 0, 25, 50, 75, or 100% for a stimulus duration of 250 to 4000 ms. Observers performed an asynchronous task at fixation throughout the experiment.

While these stimuli were being presented for the passive viewing condition, the observer was required to perform a luminance decrement task on the fixation cross. The fixation cross decremented twice in luminance for 400 ms, separated by an 800 ms inter-stimulus interval and the observer reported with a button press which decrement interval appeared darker (see (Gardner et al. 2008) for details). Decrement amplitude was adjusted according to a staircase procedure to maintain ~82% correct.

### MRI acquisition and preprocessing

Visual area mapping and cortical measurements were obtained using a multiplexed sequence on a 3 Tesla GE Discovery MR750 (GE Medical Systems) with a Nova Medical 32ch head coil. Functional images were obtained using a whole-brain T2*-weighted two-dimensional gradient-echo acquisition (FOV = 220mm, TR = 500 ms, TE = 30 ms, flip angle = 46 deg, 7 slices at multiplex 8 = 56 total slices, 2.5 mm isotropic). In addition, two whole-brain high-resolution T1-weighted 3D BRAVO sequences were acquired (FOV=240mm, flip angle=12 deg, 0.9 mm isotropic) and averaged to form a “canonical” anatomical image which was used for segmentation and surface reconstruction and session-to-session alignment. A T2*-weighted scan with the phase encoding direction reversed was collected in each session and used in combination with the FSL function TOPUP to correct for distortions due to high multiplex factors (Andersson et al. 2003). In each functional session, we also obtained a “session” anatomical image for alignment with the canonical anatomy using a T1-weighted 3D BRAVO sequence (FOV=240 mm, flip angle=12 deg, 1.2×1.2×0.9 mm). Analysis was performed using custom MATLAB software (Gardner et al. 2018a).

Session anatomies were aligned to the canonical anatomy and data were displayed on flattened cortical surfaces for visualization and for defining visual areas. Gray-matter and white-matter segmentation was performed on the canonical anatomy using FreeSurfer (Dale et al. 1999) and flattened triangulated surfaces used for displaying data. Each session anatomy, was aligned to the canonical anatomy using image-based registration (Nestares and Heeger 2000) so that the location of mapped cortical visual areas could be projected into each session’s space. All data analysis was performed in the native coordinate of the functional scan without transformation.

Cortical visual area mapping was performed using a population receptive field mapping technique (Dumoulin and Wandell 2008). Observers performed the fixation task described above while a moving-bar stimulus moved across the visual field in different directions. The measured responses were used to estimate the voxelwise population receptive field and then the eccentricity and polar angle of each receptive fields was projected onto a flattened representation of the cortical surface where visual areas were identified according to published criteria by hand (Wandell et al. 2007; Gardner et al. 2008). Each moving bar stimulus scan lasted four minutes and the same randomization sequence was repeated and averaged eight times to improve the signal-to-noise ratio. The stimulus was a full contrast 3 deg width bar spanning the entire visual field. Inside the bar a full contrast cross-hatch pattern of black and white rectangles moved continuously to minimize adaptation. Each of the four-minute scans began with a 12 s blank followed by eight 24 s cycles in which the bar swept across the entire screen in one of the eight cardinal or oblique directions. Two additional 12 s blanks occurred after the third and sixth bar sweeps to help estimate large population receptive fields. The bar swept across the visual field at 2 deg / s. The screen was crescent shaped and extended ~25 deg vertical and ~50 deg horizontal. Beyond the screen boundaries the image was blacked out to prevent artifacts from reflecting on the scanner bore. We were able to consistently map V1-hV4, V3A/B, V7 (IPS0) and hMT+ (referred to as MT, see (Huk et al. 2002; Amano et al. 2009) in all observers. Areas LO1-2, VO1-2, and IPS1-3 were not consistently identified and were therefore excluded from analysis.

Motion correction, linear trend removal, filtering, and averaging across cortical visual areas was performed to obtain a single time course for each cortical area for each observer. T2*-weighted images were motion-corrected with a rigid body alignment using standard procedures (Nestares and Heeger 2000). Scans within each session were linearly detrended, high-pass filtered with a cutoff frequency of 0.01 Hz to remove low frequency drifts, converted to percent signal change by dividing each voxel’s time course by its mean image intensity within each scan and then concatenated across scans.

Analyses of responses of cortical areas were conducted by averaging the time series of voxels whose trial-triggered response across all conditions accounted for the highest amount of variance within each retinotopically defined visual area. Specifically, we performed an event-related analysis to recover the response evoked by each trial (regardless of condition), using the following equation to model voxel responses:

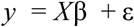

Where *y* is an *n* × 1 array representing the time-series of bold> response for n volumes from a single voxel. *X* is an *n* × *k* stimulus convolution matrix in which the first column contains a one for the volume when each trial began and zeros elsewhere. Each subsequent column is shifted downwards by one to form a Toeplitz matrix and *k* was set to 81 to model responses as occurring from the time of stimulus presentation through 40.5 s later. Each voxel is assumed to have additive gaussian noise with variance ε. By computing the least-squares estimate of the column vector β, we obtained the finite impulse response evoked by all trials, that is, the average response after a trial accounting for linear response overlap. We computed r^2^, the amount of variance accounted for by this model (Gardner et al. 2005). We then averaged the time series of the top 25 voxels per cortical area sorted by *r*^2^. While we chose this voxel selection criterion to produce high signal-to-noise estimates of each cortical areas response, our conclusions did not depend on its use. Repeating the complete analysis using either all voxels in each cortical area, the top two voxels, or all voxels weighted by their receptive field overlap with the stimulus results in a change in the signal-to-noise in the data but did not qualitatively change the results.

To examine how the hemodynamic response for each cortical area changed as a function of stimulus condition (Fig 2), we computed the finite impulse response for each condition in the passive viewing experiment. That is, we computed the finite impulse response as above, but allowed for a separate response for each of the 20 conditions in the cross set and 20 in the duration set. Our complete stimulus convolution matrix therefore had 3240 columns (81 volumes by 40 conditions), while each observer’s data consisted of at minimum 13,440 timepoints and up to 30,000 timepoints in some observers. Solving for the least squares solution results in hemodynamic response for each of the forty conditions in the experiment which we call the measured cortical response.

### Population response functions: overview

Using the measured cortical responses we then estimated the population response functions for contrast and coherence in each cortical visual area. Our model framework and measurements are available online, as a tool for experiment design and comparison with existing results (Birman and Gardner 2018). Following previous work examining the relationship between contrast or coherence and bold> response (Boynton et al. 1996, 1999; Tootell et al. 1998; Heeger et al. 2000; Rees et al. 2000; Logothetis et al. 2001; Avidan et al. 2002; Olman et al. 2004; Gardner et al. 2005) we assumed that there was a smooth functional form (linear, exponential or sigmoidal, see details below) between the contrast and coherence of the stimulus and the magnitude of neural response. For each trial, the magnitude of neural response was computed as the linear sum of the response to contrast and coherence predicted by these smooth functions and a trial onset response that was the same across all conditions (interaction terms between contrast and coherence were tested and compared against simpler models by cross-validated variance explained). The neural magnitude was used to scale the magnitude of a boxcar function of the appropriate duration exponentially scaled (see below) to account for non-linear effects of duration. The resulting time series was then convolved with a canonical hemodynamic response function estimated from the data. The parameters of the population response functions and magnitude of the trial onset response were then adjusted to best fit the event-related responses in the least squares sense through non-linear fitting routines (active-set algorithm implemented in *lsqnonlin* in MATLAB). To avoid overfitting and to compare models with different numbers of parameters, we evaluated models according to the cross-validated *r*^2^ by performing a leave-one-condition out cross-validation, using 39 of the 40 stimulus conditions to train the model while predicting on the left out condition. We proceeded with this analysis in two steps: characterizing the canonical hemodynamic response and duration effects, and then fitting the population response functions parameters.

### Population response functions: canonical hemodynamic response function and duration effects

We first fit parameters of the canonical hemodynamic response function and duration effects, ignoring the effect of contrast and coherence. To do so we fit the population response model with arbitrary scaling factors (beta weights) for each of the 40 conditions. This approach allowed us to determine the shape parameters of the hemodynamic response function and temporal non-linearity without being biased by magnitude differences across conditions.

We characterized the shape of the canonical hemodynamic response function for each observer with a difference of two gamma functions:

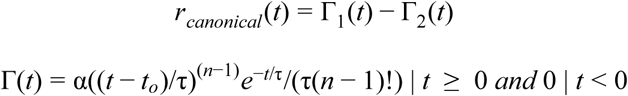

Where α is the amplitude, *t*_0_ is the time lag such that when *t* < *t*_*o*_ the function is zero, *n* and τ control the shape of the function. The parameter α was set such that the peak response to a 500 ms stimulus was 1. Thus the reported percent signal change in the population response functions are relative to a 500 ms stimulus.

We accounted for non-linear effects of temporal summation (Boynton et al. 1996) in the bold> response by allowing responses to be exponentially scaled. Small variations in duration are known to scale in an approximately linear manner (Boynton et al. 1996) whereas across large variation in stimulus durations the response to longer durations is less than expected by a linear system (Boynton et al. 2012). We are agnostic to the source of this effect, which could result from either neural adaptation (Buxton et al. 2004) or due to saturation of the bold> signal (Friston et al. 1998). We took the response of the 500 ms duration stimulus as the baseline and scaled shorter and longer responses according to the inverse ratio of the durations raised to a fit parameter δ (i.e. a 1000 ms stimulus has a ratio of 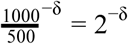). This final value corresponds to the proportion of a linear response that occurred and the boxcar of appropriate duration was scaled by this value.

Altogether we fit the parameters for two gamma functions in the canonical hemodynamic response function (α, *t*_0_, τ), the duration effect δ and 40 beta weights for stimulus conditions to the event-related responses. The canonical hemodynamic response function parameters and the duration parameter were then used in the estimation of the population response functional forms while the beta weights were discarded.

### Population response functions: functional forms

To characterize the population responses of each visual area to changes in contrast and motion coherence we fit functional forms to the underlying neural population response functions. We assumed that these population response functions would be monotonically increasing for both contrast and coherence. For contrast, we parameterized the relationship between contrast and neural response as a sigmoidal function (Naka and Rushton 1966) following previous work (Albrecht and Hamilton 1982):

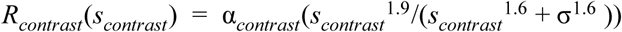

Where α is the maximum amplitude of the function and controls the shape of the function, a value near one being linear in the range we measured. We fixed the exponent parameters of the Naka-Rushton to 1.9 and 1.6 based on previous work (Boynton et al. 1999).

To avoid making assumptions about the coherence response function we assumed that the form would either be linear or a saturating nonlinearity motivated by previous work (Simoncelli and Heeger 1998; Rees et al. 2000). The saturating nonlinearity was an exponential function but can interpolate smoothly between a linear and nonlinear function.

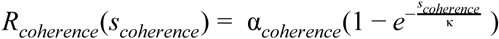

In the exponential function the parameter κ controls the shape of the function by setting the point at which the exponential function reaches 63% of its maximum and α controls the amplitude. Large values of κ combined with large values of α make this function approach linear in the range [0 1] in which the stimulus strength *s*_*coherence*_ is bounded.

To assess whether and to what extent contrast and motion coherence interact we included an additional parameter in the population response function model. The parameter β_*interaction*_ scaled the multiplicative effect of contrast and motion coherence according to the following equation:

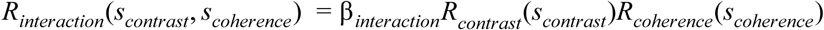

The full model of neural response was computed as the sum of the contrast and coherence response, the interaction term, and a constant stimulus onset effect *R*_*onset*_.

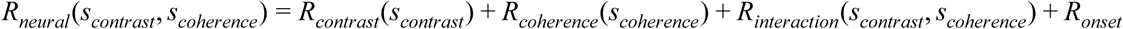

We evaluated the fit of the full model with and without the additional interaction parameter by comparing the cross-validated variance explained. We also fit an alternative interaction model in which different population response functions were allowed to fit for conditions in which only one feature changed (i.e. the first column and last row of the “grid” in Fig. 2A) compared to conditions in which both features changed (other parts of the grid in Fig. 2A).

**Figure 2.**
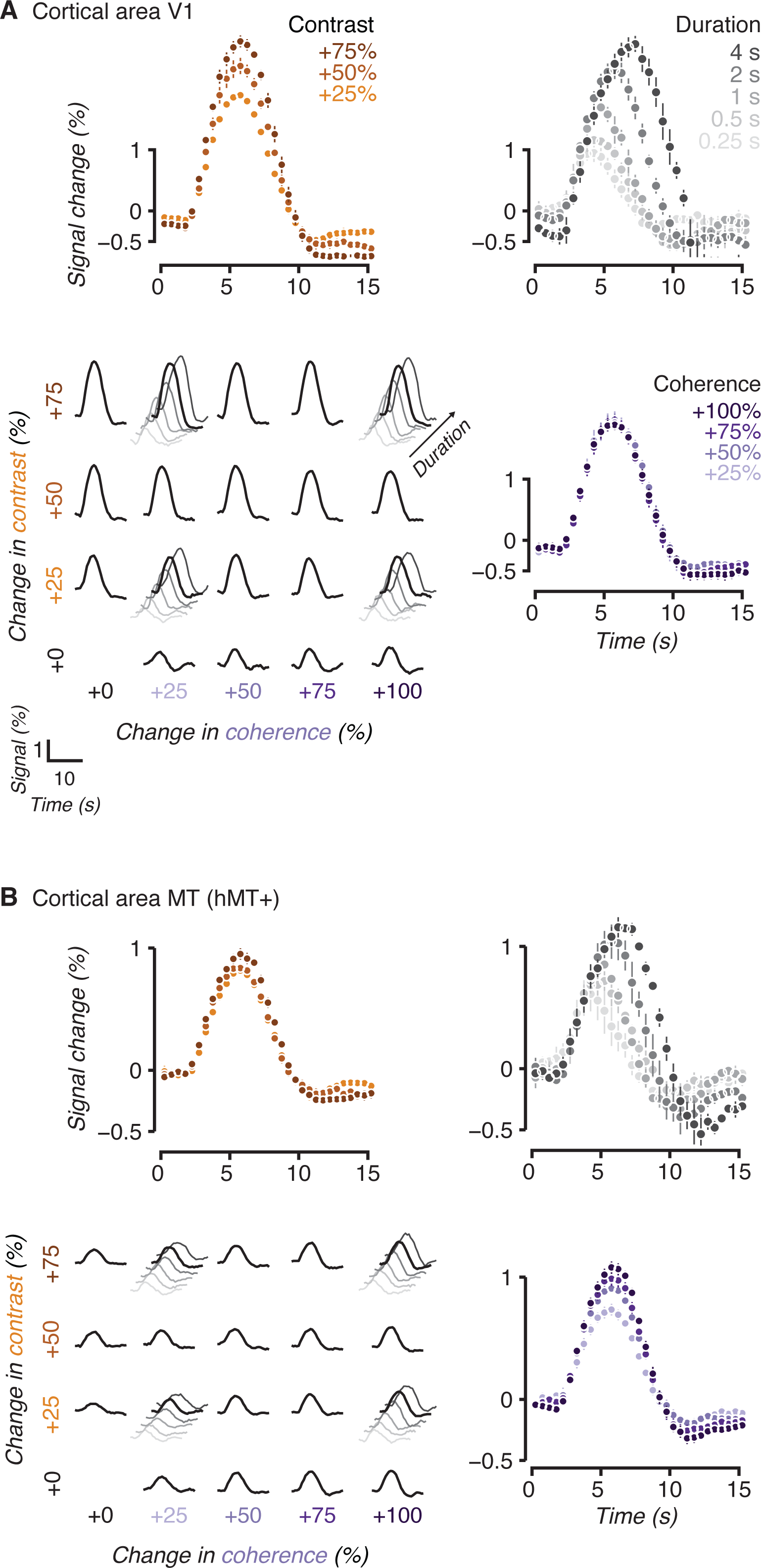
Measurements of event-related responses in cortical areas V1 and MT. (a) Cortical area V1. To obtain the individual responses shown here we performed an event-related analysis on our time series. In total we included forty conditions in the experiment: twenty consisted of a full cross of changes in contrast and/or coherence presented for 2500 ms (shown in bold in the grid in the bottom left) and twenty were a subset of the full cross conditions presented for various durations (shown in diagonal for the four conditions with additional durations recorded). We measured cortical responses to changes in contrast (top left) where each trace is averaged over changes in coherence, i.e. each response is the average of a row in the bottom left grid. We also measured responses to changes in coherence (bottom right), each trace is averaged over changes in contrast, i.e. each response is the average of a column in the grid. We made additional measurements across a large range of stimulus durations (top right) also shown in the grid. (b) As in (a) for cortical area MT (hMT+). In all panels the event-related responses are averaged across observers and error bars indicate the bootstrapped 95% confidence interval, some error bars may be hidden. Note that for visualization event-related responses are only shown out to 15 s but the analysis used a window of 40.5 s.

We fit the free parameters of the population response functions by constraining the fits on each observer’s cortical measurements (Fig. 4). To do this we computed the neural response *R*_*neural*_ and then scaled this by the boxcar of appropriate duration for each stimulus condition. The boxcar was additionally scaled according to the duration parameter. Finally we convolved this scaled boxcar with the canonical hemodynamic response resulting in a predicted hemodynamic response for each stimulus condition.

To evaluate whether the parameters we fit differed across subjects and across cortical areas we fit a linear model for each parameter. We first performed model comparison to establish whether each parameter was better explained by a model with only an intercept (parameter ~ 1), a per-subject effect (parameter ~ 1 + subject), a per-area effect (parameter ~ 1 + area), or a per-subject and per-area effect (parameter ~ 1 + subject + area). For each parameter we fit all four models (using the function *fitlme* in MATLAB) and retained the most complex model which resulted in a statistically significant improvement in prediction, assessed via partial *F*-test. For each parameter we then investigated which observers and cortical areas showed statistically significant differences relative to the mean parameter value as reported in Table. 1 and 2.

**Table 1.** Variability in parameter estimates across cortical areas. Each column reports the mean and standard deviation for one fit parameter of the population response function model averaged over observers for each cortical area. A * indicates statistically significant variability across cortical areas estimated by model comparison against a null model with only an intercept. Parameters that differ from the mean parameter value at the *p* < 0.05 threshold are shown with their p-value (probability of occurrence).

|  | Hemodynamic response |  |  | Contrast |  | Coherence |  | Onset |
| --- | --- | --- | --- | --- | --- | --- | --- | --- |
| Area | $\tau_0$ | $\tau_1^*$ | $\alpha_1$ | $\alpha_{contrast}^*$ | $\sigma^*$ | $\alpha_{coherence}$ | $\kappa$ | $R_{onset}^*$ |
| V1 | 0.51+-0.05 | 1.80+-0.42 | -0.24+-0.06 | 1.68+-1.18 | 0.35+-0.14<br>($p=0.006$) | 13.94+-32.04 | 2.11+-6.79 | 0.42+-0.24<br>($p<0.001$) |
| V2 | 0.52+-0.06 | 1.79+-0.44 | -0.24+-0.08 | 0.69+-0.57 | 0.40+-0.13 | 21.27+-27.42 | 0.40+-1.06 | 0.28+-0.16 |
| V3 | 0.52+-0.07 | 1.79+-0.59 | -0.24+-0.09 | 0.63+-0.34 | 0.43+-0.13 | 13.23+-26.11 | -0.61+-3.13 | 0.30+-0.12<br>($p=0.013$) |
| V4 | 0.52+-0.07 | 1.83+-0.93 | -0.24+-0.09 | 0.61+-0.24 | 0.47+-0.12 | 14.05+-22.42 | -1.83+-5.98 | 0.21+-0.13 |
| V3a | 0.52+-0.07 | 1.79+-0.59 | -0.24+-0.09 | 0.35+-0.26<br>($p=0.041$) | 0.48+-0.11 | 3.41+-12.46 | 1.05+-1.71 | 0.15+-0.1<br>($p=0.005$) |
| V3b | 0.52+-0.07 | 1.79+-0.59 | 0.24+-0.09 | 0.24+-0.13<br>($p=0.003$) | 0.43+-0.17 | 7.99+-16.15 | 0.29+-0.41 | 0.14+-0.06<br>($p=0.002$) |
| V7 | 0.52+-0.07 | 2.15+-0.87<br>($p=0.032$) | -0.23+-0.07 | 0.32+-0.25<br>($p=0.022$) | 0.53+-0.22 | 9.80+-14.21 | 1.11+-1.87 | 0.09+-0.07<br>($p<0.001$) |
| MT | 0.51+-0.04 | 1.81+-0.63 | -0.22+-0.09 | 0.22+-0.25<br>($p=0.001$) | 0.58+-0.32 | 6.87+-17.60 | 0.92+-1.07 | 0.24+-0.09 |

**Table 2.** Variability in parameter estimates across observers. Each column reports the mean and standard deviation for one fit parameter of the population response function model averaged over cortical areas for each observer. All of the parameters were found to have statistically significant variability between observers. Parameters that differ from the mean parameter value at the *p* < 0.05 threshold are shown with their p-value (probability of occurrence).

|  | Hemodynamic response |  |  | Contrast |  | Coherence |  | Onset |
| --- | --- | --- | --- | --- | --- | --- | --- | --- |
| Obs | $\tau_0$ | $\tau_1$ | $\alpha_1$ | $\alpha_{contrast}$ | $\sigma$ | $\alpha_{coherence}$ | $\kappa$ | $R_{onset}$ |
| 1 | 0.53+-0.03 | 2.12+-0.09<br>($p=0.041$) | 0.24+-0.01 | 0.73+-0.46 | 0.47+-0.14 | 20.64+-19.35 | 0.00+-0.00 | 0.31+-0.11<br>($p=0.013$) |
| 2 | 0.50+-0.01 | 2.09+-0.15<br>($p=0.020$) | 0.23+-0.02 | 0.23+-0.37<br>($p<0.010$) | 0.32+-0.18<br>($p=0.004$) | 5.68+-18.19 | 1.30+-1.85 | 0.28+-0.13 |
| 3 | 0.53+-0.03 | 1.80+-0.36 | 0.20+-0.03<br>($p<0.001$) | 1.31+-1.65<br>($p<0.001$) | 0.50+-0.02 | 1.77+-3.82 | 2.04+-1.55 | 0.06+-0.13<br>($p<0.001$) |
| 4 | 0.41+-0.03<br>($p<0.001$) | 1.29+-0.25<br>($p<0.001$) | 0.13+-0.03<br>($p<0.001$) | 0.56+-0.45 | 0.26+-0.20<br>($p<0.001$) | 6.81+-15.88 | 1.19+-5.85 | 0.22+-0.18 |
| 5 | 0.45+-0.02<br>($p<0.001$) | 3.09+-0.53<br>($p<0.001$) | 0.10+-0.02<br>($p<0.001$) | 0.64+-0.92 | 0.50+-0.07 | 15.81+-22.26 | 2.66+-5.01<br>($p=0.005$) | 0.29+-0.13 |
| 6 | 0.54+-0.03<br>($p=0.005$) | 2.41+-0.15<br>($p<0.001$) | 0.22+-0.02 | 0.66+-0.23 | 0.47+-0.25 | -12.93+-11.24<br>($p<0.001$) | 3.82+-7.77<br>($p<0.002$) | 0.18+-0.11 |
| 7 | 0.51+-0.03 | 1.77+-0.12 | 0.27+-0.04<br>($p<0.001$) | 0.42+-0.18 | 0.56+-0.20<br>($p=0.040$) | 3.88+-9.76 | 0.56+-0.80 | 0.12+-0.08<br>($p=0.001$) |
| 8 | 0.61+-0.05<br>($p<0.001$) | 1.47+-0.12<br>($p<0.001$) | 0.33+-0.04<br>($p<0.001$) | 0.40+-0.43 | 0.44+-0.10 | 34.68+-28.94<br>($p<0.001$) | 0.03+-0.29 | 0.33+-0.17<br>($p=0.004$) |
| 9 | 0.54+-0.03<br>($p=0.002$) | 2.05+-0.54<br>($p<0.001$) | 0.22+-0.03<br>($p=0.032$) | 0.45+-0.32 | 0.58+-0.18<br>($p=0.014$) | 12.60+-20.76 | 0.11+-0.82 | 0.32+-0.25<br>($p=0.009$) |
| 10 | 0.54+-0.03 | 1.18+-0.24<br>($p<0.001$) | 0.34+-0.05<br>($p<0.001$) | 0.52+-0.31 | 0.39+-0.16 | 7.12+-9.65 | 0.77+-1.02 | 0.20+-0.15 |
| 11 | 0.52+-0.02 | 1.02+-0.14<br>($p<0.001$) | 0.33+-0.03<br>($p<0.001$) | 0.61+-0.18 | 0.56+-0.20<br>($p=0.016$) | 28.44+-28.99<br>($p<0.001$) | 0.02+-0.02 | 0.24+-0.11 |

### Computing stimulus sensitivity

For each cortical area we computed various measures of sensitivity to contrast and motion coherence. In particular, we examined the α_*contrast*_ parameter, which control the maximum response of the Naka-Rushton function. Because in the range we measured the slopes are approximately linear and the *R*_*onset*_ term absorbs the stimulus independent response, α_*contrast*_ tracks the slope of the relationship between contrast and response and therefore is a measure of sensitivity to contrast. The parameters of the exponential form of the coherence function we used are not interpretable in isolation so instead we took the population response functions for coherence and measured their response range by performing a linear fit. We report the slope of that fit as the sensitivity to coherence.

The measurements of sensitivity which we report will be sensitive to the signal-to-noise of our measurements. This could be particularly problematic because signal magnitude and variability may depend on whether there are sinuses or large draining veins in a cortical region which are known to have large signals with high variability. Also, differences in signal-to-noise that are due to proximity to receiver coils or partial voluming effects may bias our measurements of sensitivity, particularly making comparisons across different areas problematic. In addition if variance is proportional to the mean as it is expected to be for single neurons or poisson-like processes (Softky and Koch 1993), then measures of population sensitivity would need to be scaled appropriately as response magnitude grows. We therefore examined the variability of response in each cortical visual area. First, we fit a canonical hemodynamic response function to all trials as described above. We then fit a general linear model using this canonical hemodynamic response and allowed each trial to have a separate beta-weight. That is, we found the scale factor (beta-weight) for every single trial which best fit the measured time course in the least squares sense, accounting for linear overlap across trials, for each observer for every cortical area. To avoid response variance associated with different stimulus strengths, we grouped the scale factors by condition (20 contrast and coherence; 20 duration) and computed the standard deviation. This results in 3520 measurements of standard deviation (11 observers * 8 cortical areas * 40 conditions) each of which was computed from approximately 25 trials. If the microvasculature, coil proximity, or partial voluming in different cortical areas resulted in differences in variability, or if contrast or coherence caused the variability to increase, we would expect that these measurements of standard deviation would consistently vary with those parameters. We tested for this by fitting a series of linear models in which the standard deviation depended on either an intercept alone (std ~ 1), each condition’s contrast (std ~ 1 + con), coherence (std ~ 1 + coh), cortical area (std ~ 1 + roi), or random effect of subject (std ~ 1 + (1 | subj)), and all the effects together (std ~ 1 + con + coh + roi + (1 | subj)). We also tested models in which the contrast and coherence effects could differ by roi (std ~ 1 + con*roi; std ~ 1 + coh*roi). We performed model comparison by testing for improvement over the intercept-only model via partial *F*-test.

## Results

### Measuring cortical responses to contrast and motion coherence

We characterized human cortical responses to changes in contrast and motion coherence of patches of dynamic random-dot stimuli by measuring bold> responses while observers passively viewed two patches of moving dots (Fig. 1). Each scan began with 30 s of baseline stimulus presentation (0% coherence, 25% contrast) after which trials consisting of brief increments (0.25 - 4 s) in either or both coherence and contrast before returning back to baseline for a random length inter-trial interval (2 - 11 s) (see Methods for full details). In total observers were shown forty conditions: twenty consisted of combinations of changes in contrast (+0, +25, +50, and +75%) and changes in motion coherence (+0, +25, +50, +75, and +100%) for 2500 ms each, the remaining twenty were a subset of these combinations combined with variable stimulus durations (250, 500, 1000, 2000, and 4000 ms). To minimize task-dependent effects and maintain a consistent level of engagement observers performed an independent fixation task during viewing. We computed hemodynamic responses to each stimulus condition for each observer using an event-related analysis for retinotopically defined visual areas V1, V2, V3, hV4, V3A, V3B, V7, and MT. We begin by describing responses in visual areas V1 (Fig. 2A) and MT (Fig. 2B), as they are well-known to be sensitive to contrast (Tootell et al. 1995, 1998; Boynton et al. 1996, 1999; Logothetis et al. 2001; Avidan et al. 2002; Olman et al. 2004; Gardner et al. 2005) and motion coherence (Britten et al. 1993; Simoncelli and Heeger 1998; Rees et al. 2000; Händel et al. 2007).

We observed clear parametric sensitivity to increases in contrast in V1 but weaker sensitivity in cortical area MT. Our measurements in V1 confirm previous results (Tootell et al. 1995, 1998; Logothetis et al. 2001; Gardner et al. 2005). The contrast sensitivity of V1 can be appreciated as monotonically increasing response magnitudes for higher levels of contrast increments (top left orange traces, Fig. 2A). These traces are for a stimulus duration of 2.5 s collapsing across motion coherence increments, i.e. averaging each row in the full response grid. While MT was also sensitive to increments of contrast, the monotonic increase appeared less pronounced compared to V1 (top left orange traces, Fig. 2B), consistent with other reports that have noted MT as having near maximal responses to small changes to contrast (Sclar et al. 1990; Tootell et al. 1995).

For motion coherence, we found the opposite pattern: MT was much more sensitive to increments in motion coherence compared with V1. MT showed clear monotonic increasing responses with increasing motion coherence (bottom right purple traces, Fig. 2B). These traces are again for a stimulus duration of 2.5 s averaged over contrast increments, i.e. collapsing each column in the full response grids. In V1 there was little difference in response amplitude as a function of motion coherence, i.e. weak sensitivity to coherence (bottom right purple traces Fig. 2A).

While V1 showed little parametric sensitivity to difference in coherence and MT little sensitivity to difference in contrast, both show a large response to the smallest increment of these parameters. This consistent trial-by-trial response, which we call the stimulus-onset response, appears unrelated to our parametric manipulations. For example, despite showing little sensitivity to different levels of coherence all of the responses for V1, including the one induced by the least change in coherence (+25%), induced a large response relative to the baseline (purple traces, Fig. 2A). Similarly, for MT and contrast as can be appreciated by noting that increasing contrast by 25% (orange traces, Fig. 2b) resulted in a large response. Part of this apparently large response is due to the fact that these responses for contrast or coherence are averaged over changes in the other parameter. That is, increases in contrast are shown averaged over coherence and vice-versa. However this is not the complete story as can be appreciated by examining the grid of responses to each parameter separately (small bold black traces in grid, Fig. 2A and B). V1 can be seen to respond to a small change in coherence (+25, along horizontal) when there is no change (+0, along vertical) in contrast and vice-versa for MT. These relatively large responses, to a feature each area is not strongly sensitive to, suggests that there is a response to stimulus onset regardless of condition.

Motion visibility is also adjusted by reducing the duration of stimuli, often in conjunction with reduced contrast and coherence. Along with the measurements described above, for which the stimulus duration was 2.5 s, we tested a large array of different durations from 0.25 to 4 s. As expected of an approximately linear system (Boynton et al. 2012) we observed that responses scaled with stimulus duration in both cortical areas V1 and MT (top right grey traces, Fig. 2A,B).

Across the rest of the visual areas that we were able to retinotopically define in all subjects (V2, V3, hV4, V3A, V3B, and V7) we found similar parametric sensitivity to contrast, motion coherence and stimulus duration (Fig. 3). In general, and in concordance with previous reports (Avidan et al. 2002) we found less parametric sensitivity to changes in contrast for visual areas higher up in the visual hierarchy in the range we measured (+25 to +75% contrast). Sensitivity to coherence was observed in a number of the visual areas, although MT and to a lesser extent V3A were the clear stand-outs in showing monotonically increasing responses to this parameter. These observations will be quantified below.

**Figure 3.**
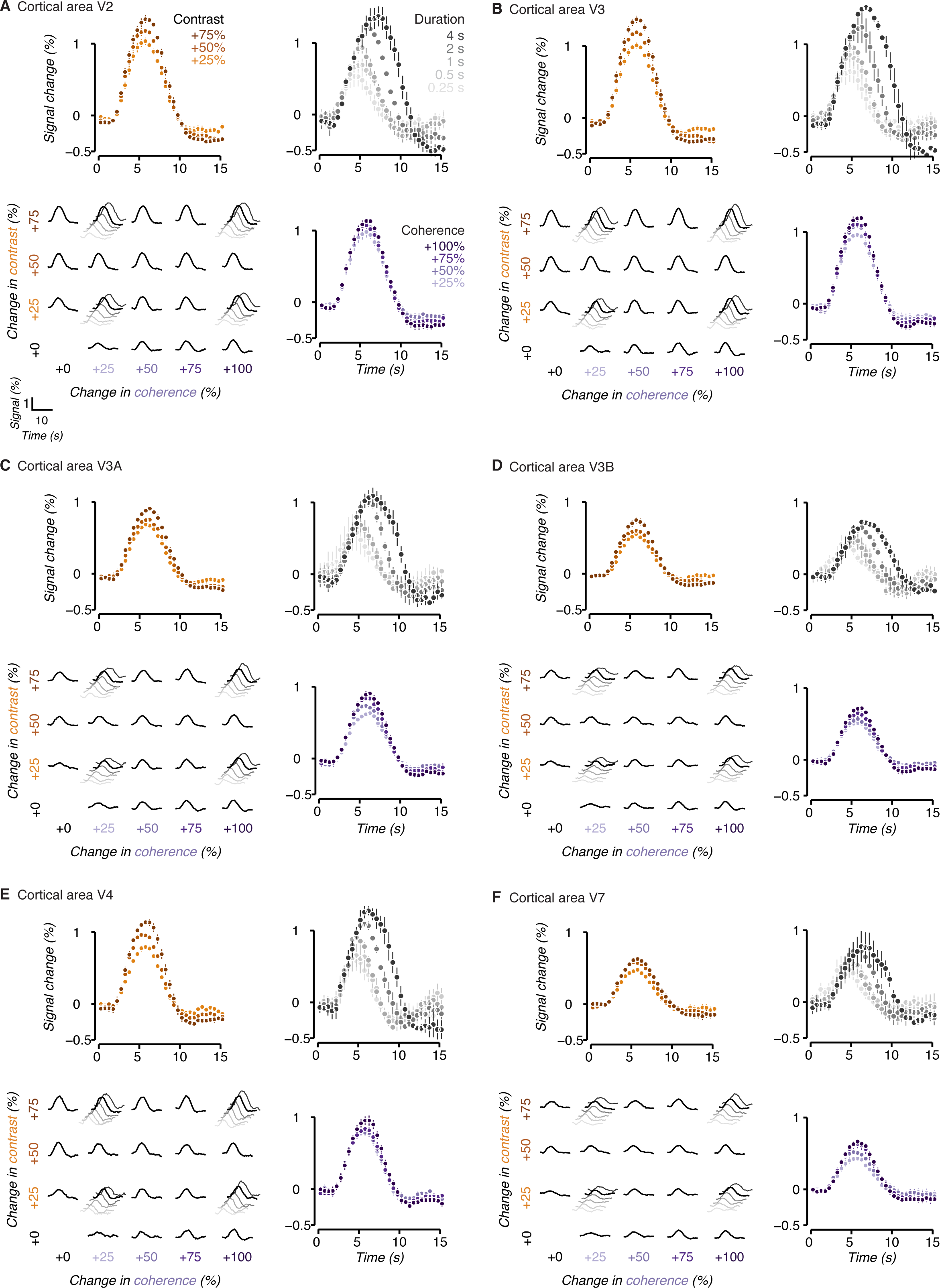
Measurements of event-related responses in cortical areas V2-V7. (a-f) Conventions are the same as in Figure 2.

### Fitting population response functions to cortical responses

To quantify the parametric sensitivity to contrast and coherence of each visual area we fit the event-related responses with a population response model using idealized functional forms for the relationship between contrast and coherence and neural response (Fig. 4). Based on previous work we expected that the population response to contrast would be a sigmoidal function (Albrecht and Hamilton 1982; Sclar et al. 1990; Boynton et al. 1999) with the form of a Naka-Rushton equation (Fig. 4B, orange curve) (Naka and Rushton 1966). To avoid overfitting, we fixed the exponents in the equation based on previous work (Boynton et al. 1999) and only allowed σ and α_*contrast*_ to vary. For motion coherence, we allowed for a functional form that can smoothly interpolate between linear (Britten et al. 1992, 1993; Simoncelli and Heeger 1998; Rees et al. 2000) and a saturating exponential (Fig 4B, purple curve). Finally, we included an onset term to capture the portion of response that did not vary across all conditions which presumably reflects stimulus onset and not parametric variation of stimulus parameters.

**Figure 4.**
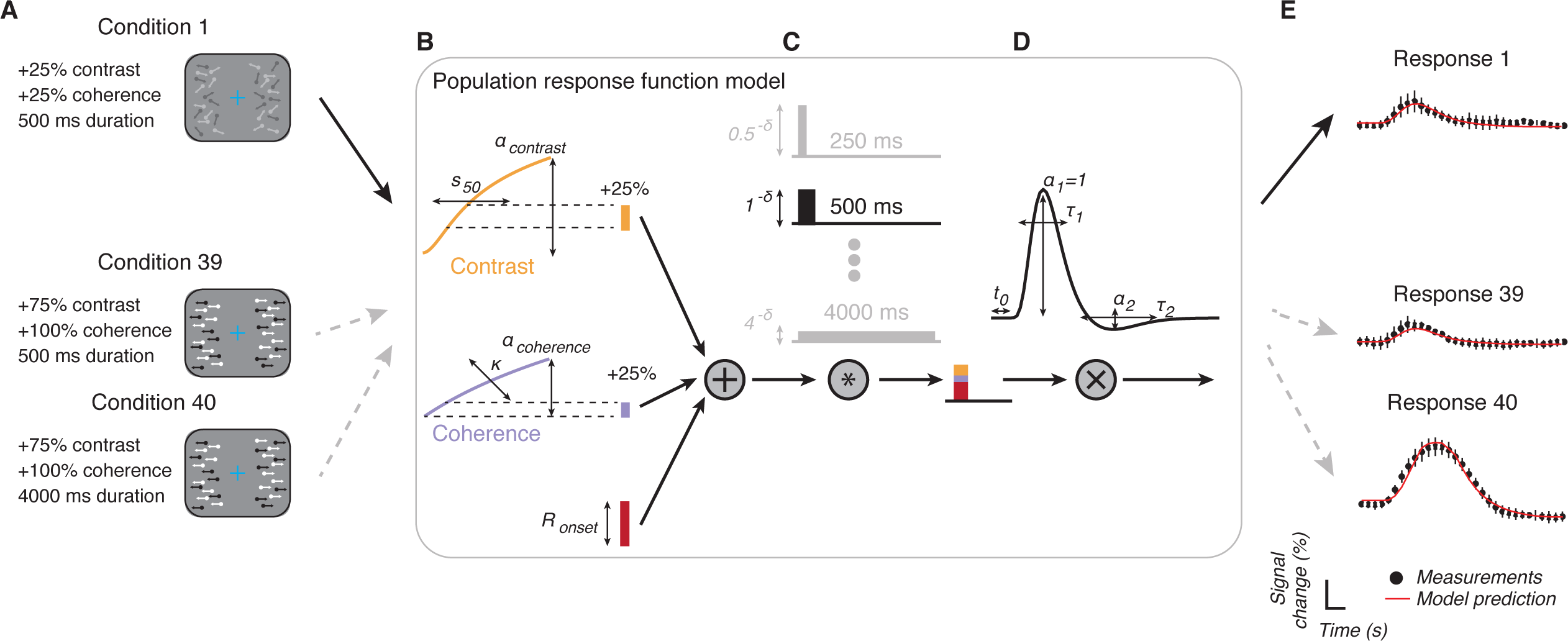
Population response function model. (a) Each condition in the experiment was defined by three parameters: the increment in contrast above baseline (+0, +25, +50, or +75%), the increment in coherence above baseline (+0, +25, +50, +75, +100%), and the stimulus duration (250, 500, 1000, 2000, or 4000 ms). As an example we use condition 1 to demonstrate the model. (b) To estimate the response to each feature within a condition we first find the change in response due to the corresponding change in stimulus intensity according to the population response functions. For contrast the population response function is a Naka-Rushton with two free parameters: α_*contrast*_ controlling the amplitude and σ the shape. For coherence the response function was a saturating nonlinearity with two free parameters: α_*coherence*_ controlling the amplitude and κ the shape. We added the resulting change in response together (while testing for interaction effects, see Methods) and included an onset parameter to account for stimulus response that did not vary parametrically with the stimulus features. (c) The total response, including onset, was used to scale a boxcar function whose length matched the stimulus duration. The boxcar was additionally scaled by a parameter to account for the nonlinear effect of stimulus duration. (d) The resulting boxcar was convolved with a canonical hemodynamic response function fit separately for each observer. (e) The model outputs a prediction for each condition about the expected event-related response (red lines). The parameters within the population response function model were then optimized to minimize the sum of squared errors between the data (black markers) and the model responses.

To predict the bold> response from the modeled contrast and coherence response functions, we employed a linear-systems approach (Heeger et al. 2000; Rees et al. 2000; Logothetis et al. 2001). To account for different durations of stimuli, we multiplied the response magnitude predicted by the onset, contrast, and coherence functions with a boxcar function of appropriate length (Fig. 4C). As it is known that brief stimuli evoke response larger than expected by linearity (Boynton et al. 1996, 2012), we also scaled the boxcar magnitude with an exponential that accounted for this non-linearity in response. This scaled boxcar was then convolved with a hemodynamic response function (Fig. 4D) whose parameters were adjusted to best fit the event-related responses across all conditions (Fig. 4E). All together, we fit the model parameters for the contrast function (α_*contrast*_, σ), coherence function (α_*coherence*_, κ) and temporal effects (δ, *R*_*onset*_), and the parameters for the hemodynamic response function (*t*_0_, τ_0_, *t*_1_, τ_1_, α_1_) for each observer for each visual area by minimizing the sum of least squares between the output of the model and the event-related responses for each of the 40 conditions.

We report the main fit parameters of the hemodynamic response function and population response function model across cortical areas (Table 1) and observers (Table 2). We assessed whether between-observer variability existed by fitting a linear model predicting each parameter with observers as categorical predictors and used the same procedure to assess for within-observer variability across cortical areas (see Methods). We found that there was statistically significant between-observer variability across all of the parameters but only significant variability within-observer (i.e. across cortical areas) for the shape parameter of the hemodynamic response τ _1_, the magnitude and shape parameters of the contrast response function α_*contrast*_ and σ, the parameters of the coherence response function α_*coherence*_ and κ, and the onset parameter *R*_*onset*_ (significance established by a partial *F*-test comparing linear regression models with and without each group of additional parameters at the *p* = 0.05 threshold). Note that the κ and α_*coherence*_ parameters which together control both the shape and magnitude of the coherence response are hard to interpret in isolation.

The population response model was able to capture the majority of variance in each observer’s event-related responses and a significant portion of this explained variance was accounted for by the population response functions. We assessed variance explained as the squared correlation between the model predictions and the actual event-related responses for held-out conditions. For V1, *r*^2^ CI [0.63 0.75]; V2, *r*^2^ = 0.63 95% CI [0.58 0.68]; V3, *r*^2^ = 0.62 95% CI [0.56 0.68]; hV4, *r*^2^ = 0.69 95% *r*^2^= 0.44 95% CI [0.35 0.53]; V3A, *r*^2^ = 0.42 95% CI [0.35 0.50]; V3B, *r*^2^ = 0.38 95% CI [0.31 0.46]; V7, *r*^2^ = 0.32 95% CI [0.24 0.40]; MT, *r*^2^ =0.49 95% CI [0.43 0.56]. Part of the variance accounted for by the model is simply due to the stimulus-onset term and hemodynamic response, but the population response functions also captured significant variance. We assessed this by comparing our results to a model fit to the same measurements but where the condition labels were permuted. This corresponds to keeping the variance explained by stimulus onset and the hemodynamic response but randomizes the relationship between condition and response. We repeated this permutation test procedure 100 times per observer and cortical area. On average across observers and areas the variance explained by fitting to the measured dataset (average cross-validated *r*^2^ = 0.508) exceeded the variance explained in the permuted dataset (average cross-validated *r*^2^ = 0.340) with *p* < 0.001, Δ*r*^2^ = 0.164, 95% CI [0.162 0.165].

Across cortical visual areas the model captured the response to changes in contrast and motion coherence as well as the amplitude effects due to duration. To visualize the fit of the population model to each variable we scaled the canonical hemodynamic response function for each observer to fit the event-related responses in the conditions with either no change in contrast or no change in coherence. This results in a single scaling factor for each of these conditions (circles, Fig 5) which we compared to the model predictions (lines, Fig 5). Examination of the magnitude of the population model fit to the event-related response peaks for changes in contrast (orange curves, Fig 5A) and coherence (blue curves) shows good correspondence. This is particularly notable given that the model is fit across all conditions containing different response lengths, as well as combinations of contrast and coherence changes, while the displayed data are for changes in contrast and coherence in isolation. This visualization displays a model fit to all the data, i.e. not on held-out data, but with similar explained variance to the cross-validated model (difference between cross-validated and full fit, Δ*r*^2^ = 0.005, 95% CI [0.004 0.006]). The population response functions echoed the qualitative results described above for the event-related responses: V1-hV4 showed strong response to contrast with relatively weak response to coherence. Only MT showed stronger response to motion coherence than to contrast. Moreover, the amplitude of responses as a function of duration (Fig 5B) were similarly well captured by the population response model. As noted earlier the amplitude of responses due to doubling in duration do not appear to scale in a linear manner.

The form of the contrast response function has been extensively studied (Albrecht and Hamilton 1982; Sclar et al. 1990; Boynton et al. 1999) while the motion coherence response function has received much less attention. Single-unit studies have found a linear response function, whereas bold> measurements in humans have found some non-linearity of response, particularly outside of MT (Rees et al. 2000). We therefore tested for nonlinearity in the population response functions to motion coherence and found that responses were generally best characterized as linear, with a small deviation from linearity for MT. We quantified this comparison as the difference in cross-validated variance explained between the saturating exponential and a linear form for the coherence response function. In MT we found a small difference in favor of the nonlinear model Δ*r*^2^ = 0.004 (95% CI [0.001 0.007]) while all other cortical areas’ confidence intervals overlapped with zero. This difference is visible as the saturation of the MT coherence response to large changes in coherence (Fig. 2B and Fig. 5A, MT).

**Figure 5.**
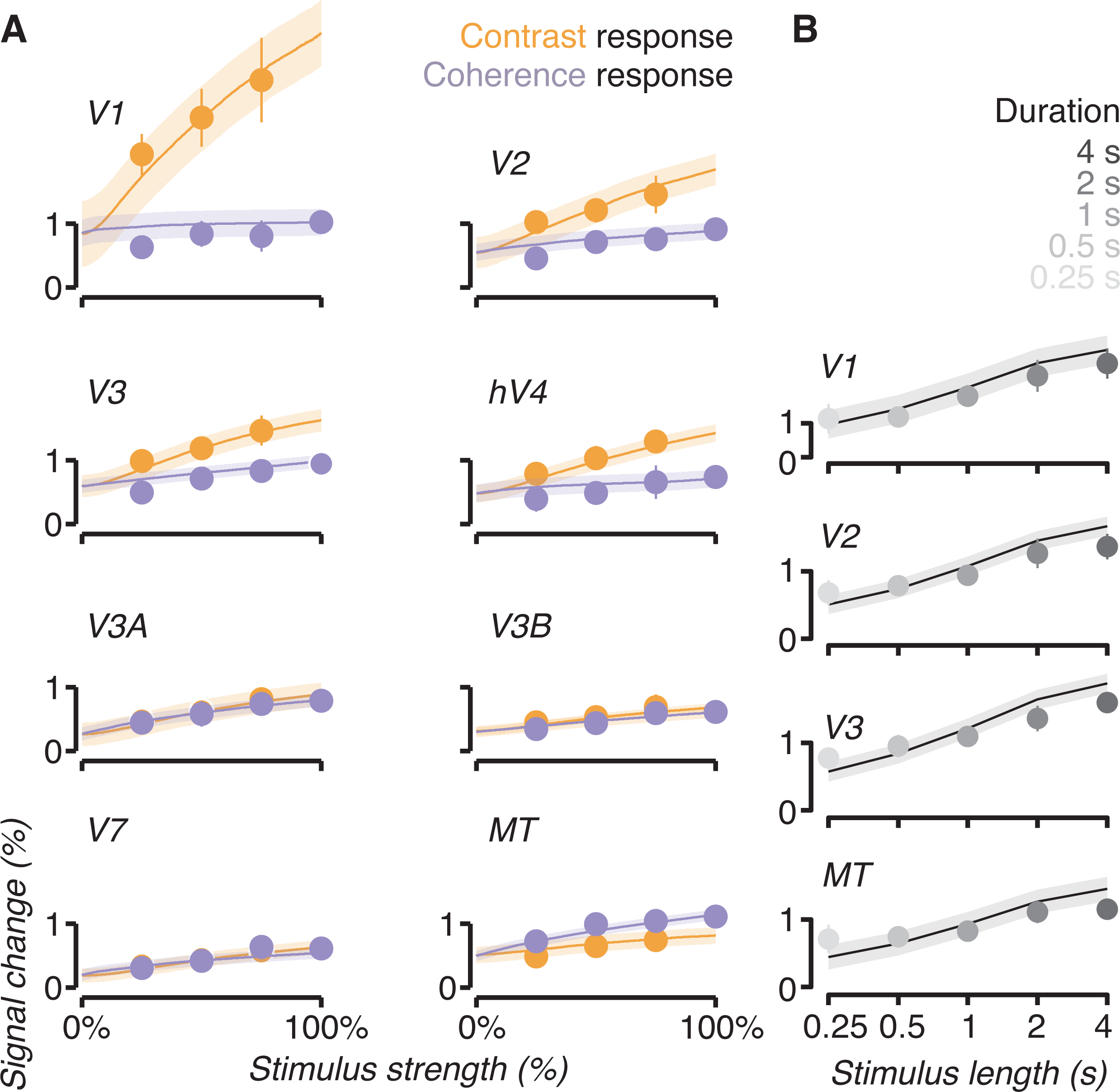
Population response functions. (a) The population response functions fit to each cortical area V1-MT (hMT+) are shown compared to the magnitude of the event-related response for the conditions in which only one feature changed. These correspond to the conditions in the first column and last row of each event-related response grid in Figure. 2 and 3. To make the functions comparable to the data in an easy to interpret space we reduced each event-related response to a single magnitude value which was obtained by finding the linear scaling of the canonical hemodynamic response to that condition. The model outputs predictions for all forty conditions but we are only showing the subset where either contrast or coherence changed alone. Note that the predictions here are not out of sample (i.e. these are not the cross-validation results) but we show the full fit to better visualize the response functions. (b) As in (a) but for the variable duration conditions in which contrast and coherence changed maximally (+75% contrast, +100% coherence). In all plots markers indicate the average across observers and error bars the bootstrapped 95% confidence interval.

While population responses to each motion feature could interact, i.e. a change in contrast might influence the response to a change in coherence or vice versa, we found no evidence for this. We tested for interactions by adding an additional beta weight to the model accounting for the effect of multiplicative changes in contrast and coherence (see Methods section *Population responses: functional forms*). Including this term reduced the cross-validated variance explained by on average −6.67% (95% CI [−13.42, 0.08]) across cortical areas, suggesting overfitting compared to the no-interaction model. One observer’s data was particularly strongly overfit. Removing that observer resulted in an average reduction in variance explained of −0.08% (95% CI [-0.25 0.09]) and for individual areas, V1: −0.18%, V2: −0.16%, V3: −0.17%, hV4: −0.13%, V3A: −0.07%, V3B: −0.14%, V7: −0.07%, MT: −0.11%.

Visual inspection of the response grids (bottom left panels in Fig. 2 and 3) suggest an alternative kind of interaction in which the response to contrast and coherence might be stronger in the absence of the other feature changing. Take for example the response to contrast compared to coherence in V1. The contrast response in V1 is so much larger than the response to coherence that it’s possible it “washes out” any visible effect due to coherence. To test for this possibility we fit a model with different population response functions for conditions in which only a single feature changed vs. when both features changed. We found that these models were also not statistically better than the simplest model with no interactions: average reduction in cross-validated variance explained -5.34% 95% CI [-9.13, -1.56] and without the overfit observer −0.20% 95% CI [-0.31, −0.08]. Although statistically the models were similar in our dataset we did find that in the interaction model the population response functions to contrast had a higher maximal response when the coherence was not simultaneously changed, but the reverse was not true. On average across subjects and cortical areas we found an increase in sensitivity of about 50% in the contrast response when no simultaneous change in coherence occurred (average parameter change 1.68 95% CI [0.58, 2.78], significantly different from zero as assessed by bootstrap over observers, *p* = 0.007).

The population response model fits (Fig. 5) replicate earlier reports showing that contrast responses have a smaller dynamic range and saturate more quickly in higher visual cortical areas (Avidan et al. 2002), and add the finding that coherence sensitivity peaks in MT. To assess this we plotted the maximum of the contrast response function (the α_*contrast*_ parameter) against the linear slope of the coherence response function (the response range measured as the slope of a linear fit, see Methods) for each cortical area (Fig 6A). As expected we found stronger sensitivity to motion coherence in V3a and MT compared to area V1 (Zeki et al. 1991; Watson et al. 1993; Dupont et al. 1994; Tootell et al. 1995). The difference in coherence sensitivity between V3A and V1 was 0.167, *p* < 0.001 and between MT and V1 0.251, *p* < 0.001. But we also observed significant sensitivity to changes in coherence in all regions measured (Fig. 6B): V1 = 0.12, V2 = 0.19, V3 = 0.18, hV4 = 0.13, V3A = 0.25, V3B = 0.15, V7 = 0.22, MT = 0.36, slopes in % signal change / unit coherence, all *p* < 0.001 assessed by bootstrap across observers. All cortical visual areas showed statistically significant parametric sensitivity to changes in contrast (Fig. 6C) assessed as a non-zero α_*contrast*_ parameter by bootstrap across observers, all *p* < 0.001 except MT, *p* = 0.002. The maximum contrast response dropped quickly for regions higher in the visual hierarchy (V1 = 2.00, V2 = 0.87, V3 = 0.68, hV4 = 0.63, V3A = 0.35, V3B = 0.24, V7 = 0.33, MT = 0.20, units in % signal change / unit contrast).

**Figure 6.**
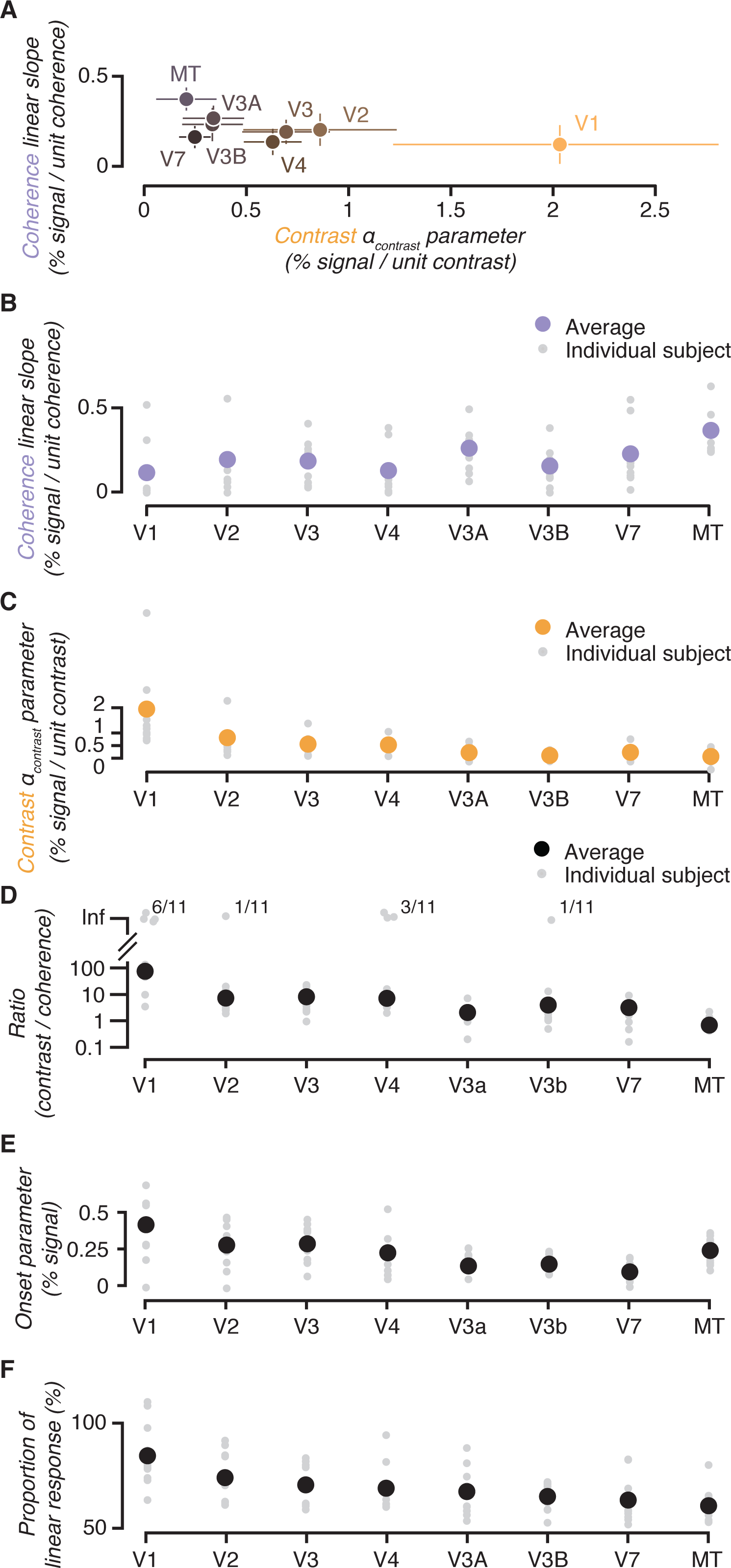
Cortical sensitivity to contrast and motion coherence. (a) To obtain a qualitative estimate of cortical sensitivity to each motion visibility feature across the cortical visual areas we plotted the α_*contrast*_ parameter from the Naka-Rushton function against the slope of a linear fit of the coherence functions. (b) The slope of the coherence functions fits as in (a) replotted with individual subjects. (c) The α_*contrast*_ parameter as in (a) replotted with individual subjects shown for each cortical area. (d) Because the amplitude parameters can be sensitive to the signal-to-noise ratio of the measurement in each cortical area their absolute magnitude is difficult to interpret. To remove any bias due to this we plot the ratio of the contrast and coherence parameters. Note that for some subjects the slope of the coherence response was near zero in some cortical areas, we note these as a ratio of infinity (Inf). The means are calculated excluding infinite values. (e) The stimulus-onset response parameter_*Ronset*_ indexes the portion of the response that was not parametrically modulated by contrast or coherence. (f) For each doubling in stimulus duration the proportion of response increase is shown by cortical area where 100% would indicate that responses increased linearly with duration. In all panels markers indicate the mean and error bars the bootstrapped 95% confidence interval. Error bars are omitted in panels (b-e) for visualization.

Although we fit a Naka-Rushton function to the contrast response our measurements were limited to only a few points (no change in contrast, +25, +50, and +75%). This meant that the data did not strongly constrain a sigmoidal fit. We assessed whether in our dataset the results would be equally well fit by a linear model and found that this was the case for all areas except V7, with an average improvement of 0.32% in cross-validated variance explained. Therefore, the α_*contrast*_ parameter which fits the maximal response to contrast in each region tracks the slope of the relationship between contrast and response and can therefore be used as a measure of the sensitivity to contrast, in the range of contrasts we measured. The linear model’s improvement in variance explained for individual areas were: V1 0.006, 95% CI [0.001 0.010]; V2 0.007, 95% CI [0.001 0.013]; V3 0.005, 95% CI [0.000 0.009]; V4 0.003, 95% CI [0.001 0.004]; V3A 0.002, 95% CI [0.001 0.004]; V3B 0.002, 95% CI [0.000 0.003]; V7 −0.000, 95% CI [-0.001 0.001]; MT 0.002, 95% CI [0.001 0.003].

We found that the variability in our measurements did not differ significantly across different cortical areas or according to the stimulus strength. We performed this analysis to test whether various nuisance variables could have altered our measurements, e.g. proximity to the coils and partial voluming might affect signal-to-noise in different cortical areas, or the variability in our measurements might increase with response magnitude as contrast and coherence cause populations of neurons to be more active. To do this we estimated the response magnitude of every trial and grouped these by condition and cortical area, then fit a series of linear models to see whether variability differed. We found that none of the additional variables improved the model fit over the intercept-only model at the *p* = 0.05 significance threshold. Importantly, the model which allowed separate values for each cortical area did not improve the model fit, suggesting that response variability did not significantly differ between cortical areas (mean cortical area standard deviation was 1.50 percent signal change; V1 1.75, 95% CI [1.33 2.17]; V2 1.08, 95% CI [0.91 1.25]; V3 1.28, 95% CI [1.06 1.49]; V4 1.37, 95% CI [1.03 1.72]; V3a 5.71, 95% CI [1.28 10.15]; V3b 1.14, 95% CI [0.96 1.31]; V7 1.32, 95% CI [0.97 1.66]; MT 1.30, 95% CI [0.95 1.66]). In addition we found that there was no statistically significant change in variability as contrast or coherence increased (even when separate slopes were allowed for different cortical areas), suggesting that noise in our measurements was additive, i.e. did not increase with increasing response magnitude. Fitting the model with a slope for contrast and coherence (shared across areas) results in a slope of -1.23 percent signal change per unit contrast, *t(2557) = -1.27, p = 0.21*, and a slope of −0.58 percent signal change per unit coherence, *t(2557) = −0.78, p = 0.44*.

Although our measurements do not suggest that any bias is introduced by potential signal-to-noise differences across areas, we computed the ratio of the contrast and coherence slope parameters as an additional unbiased analysis (Fig. 6D). This ratio allows for between region comparison of the sensitivity to contrast and coherence because the ratio reports how sensitive each region is to contrast compared to coherence and not overall sensitivity. That is, the ratio should be invariant to differences in signal-to-noise, under the assumption that contrast and coherence sensitivity are equally affected. In line with our previous results we found that V1 has a ratio of contrast to coherence sensitivity that is at least an order of magnitude more than the other areas. In addition MT was found to have a ratio near 1 and lower than the other cortical areas, reflecting its stronger relative sensitivity to coherence.

We found that the portion of the bold> response that did not vary parametrically with contrast or coherence, the stimulus-onset response *R*_*onset*_, did vary across cortical areas (Fig. 6E and Table 1). On average the onset response was 0.23 percent signal change across observers and cortical areas. The stimulus-onset response in V1 and V3 were larger than average at 0.42 percent signal change, 95% CI [0.28, 0.55], while areas V3A, V3B, and V7 were smaller than average, 0.14, 95% CI [0.07, 0.20]; 0.15, 95% CI 0.11, 0.18]; 0.09, 95% CI [0.06, 0,13], respectively. The other cortical areas’ onset effects were V2 0.28, 95% CI [0.19, 0.37]; V3 0.22, 95% CI [0.22 0.36]; V4 0.23, 95% CI [0.15, 0.31]; MT 0.24, 95% CI [0.19, 0.29].

Finally, we found that the effect of increasing stimulus duration was not consistent across cortical areas (Fig. 6F). We found that early visual cortex, V1 in particular, was significantly more sensitive to changes in duration than later visual areas, especially MT. The effect of a doubling in duration on the population response, as a proportion of that expected from a linear model, was 68.56%. On average across subjects we found that V1 and MT differed significantly from this value, as assessed by bootstrap. We found that the effect of a doubling in duration in V1 was 83% of the linear model, 95% CI [76.00, 92.27], suggesting that V1 is more sensitive to stimulus duration. By contrast in MT the effect was only 62% of the linear model, 95% CI [57.06, 66.08], suggesting that MT may have a more transient response. The effects in other areas were not significantly different from the average: V2 73%, 95% CI [67.20, 79.70]; V3 70%, 95% CI [65.05, 74.95]; V4 70%, 95% CI [64.21, 76.59]; V3A 67%, 95% CI [61.24, 73.31]; V3B 64%, 95% CI [60.45, 67.64]; V7 64%, 95% CI [58.07, 70.02]

## Discussion

We have developed a quantitative framework for modeling human cortical response to motion visibility as parameterized by image contrast, motion coherence, and duration. Our results provide a comprehensive view of the variability in cortical sensitivity to these features, each of which is a critical component of visual stimuli often manipulated in experiments designed to understand visual perception and decision-making. Our measurements show that the range of responses to different levels of contrast was larger in early visual cortex, especially V1, and the range of responses for coherence larger in V3A and MT (hMT+). Nonetheless, a change in either feature caused a cortical response in all the retinotopic areas we mapped. Our results weigh on various other findings in the literature: the precise shape of population response functions, the influence of stimulus duration on cortical signals, and whether or not sensory representations for different features interact. Finally, we believe that this parameterized model, and parametric models in general, suggest mechanisms for the read out of sensory representations from population responses and have therefore made our measurements and framework available online as a resource (see Methods).

We studied changes in contrast, coherence, and duration to measure human cortical response within a range where typical human perceptual experiments are performed. One choice we made was to measure contrast from a relatively high baseline. Because the contrast response function is known to adapt to the current background stimulus without altering the form of parametric modulation (Ohzawa et al. 1982, 1985; Sclar et al. 1985, 1989; Gardner et al. 2005) the relative sensitivities we measured should hold at other baselines. With this design we were also able to show that sensitivity to changes in contrast and coherence do not interact. The interaction analysis would be impossible in stimuli where the dots appear from a black or grey background such that both contrast and coherence always change together (Britten et al. 1993; Rees et al. 2000). When designing the dot motion stimulus we also had to ensure that there were sufficient dots and a large enough aperture to be clearly visible and generate a reliable coherence response. At low dot densities the response to changes in coherence are negligible (Smith et al. 2006) and small aperture sizes can cause changes in coherence to result in decrements in response (Becker et al. 2008; Costagli et al. 2014; Ajina et al. 2015). By creating a large stimulus with high density we guaranteed that our dot motion would blanket the population receptive fields of all the cortical areas measured.

We set our stimulus to move at a constant rate of 6 deg / s, within the peak range of speed tuning in visual cortex, and used a dot stimulus rather than gratings to avoid having spatial frequency tuning affect our measurements. Although individual V1 and MT neurons in the macaque differ greatly in their speed tuning the average tuning of the population is quite similar and centered near 6 deg / s with ranges that extend far above and below that (Priebe et al. 2006). Measurements of speed tuning in humans evidence broad variability across all of visual cortex but our chosen speed is within the peak range (Singh et al. 2000; Hammett et al. 2013). One common concern with speed tuning in gratings is that spatial frequency tuning differs across cortex and directly impacts sensitivity to other stimulus properties, such as image contrast (Priebe et al. 2003, 2006). We used a random dot stimulus with a wide range of spatial frequency components rather than gratings with a specific spatial frequency to avoid this confound. In principle our stimulus drives neurons with a wide range of tunings and by averaging over voxels in each cortical area we reduce the impact of columnar and other local microstructure in each area (Liu and Newsome 2002; Sun et al. 2007).

We reported here several parameters which together defined the population response functions, but which of these represents a good measure of the sensitivity of a region? We use the term sensitivity to capture parametric differences in response magnitude with differences in contrast or coherence. Thus, an area with high contrast or coherence sensitivity is one in which the response to the lowest and highest values of these parameters evoke the largest difference in response (see the methods for how the reported parameters correspond to this). This measure can be used to compare to human behavioral contrast or coherence discrimination performance since signal detection theory predicts that perceptual sensitivity, d’, is directly proportional to this difference (Tolhurst et al. 1983; Newsome et al. 1989; Boynton et al. 1999; Zenger-Landolt and Heeger 2003; Pestilli et al. 2011). However, d’ is also inversely proportional to the standard deviation of response which could vary across different areas, particularly for measurement related reasons that would therefore distort our measures of sensitivity. But our analysis of the variability of response across different areas did not find differences, thus suggesting that our measures are an accurate reflection of contrast and coherence sensitivity. Moreover, we used a selection criteria to analyze a subset of voxels that show consistent trial-to-trial responses to reduce the effect of measurement noise but our parametrization will still be sensitive to any noise that remains.

Response variability might also change with response amplitude as it is known to do for single-unit responses. Although occasionally single neurons can be found that match perception (Britten et al. 1992), groups of neurons (Tolhurst et al. 1983) or larger populations (Zohary et al. 1994; Averbeck et al. 2006), depending on the correlation structure in the population, are likely to more closely reflect perceptual reports. Supporting the idea that populations are used for perceptual readout is evidence from human work where at the coarse resolution of the bold> signal, which pools over large numbers of neurons, cortical measurements closely track perception under an assumption of additive noise (Boynton et al. 1999; Sapir et al. 2005; Pestilli et al. 2011; Hara and Gardner 2014). In line with this the variance of population responses measured with voltage-sensitive dyes do not change with magnitude of response in V1, i.e. they are additive (Chen et al. 2006). Our own measurements support the hypothesis that populations are subject to additive noise: we found that as contrast and coherence increased and caused larger magnitudes of response we found no evidence that trial-by-trial variability changed. Together our data and previous results suggest that measures of the slope in the bold> signal population response function are indeed measures of sensitivity and leaves us with a testable prediction: if parameters measure sensitivity (i.e. signal-to-noise ratio) then they should be relatable to human perception under additive noise but not noise which scales with response magnitude.

We observed a saturation of the cortical response to motion coherence which differs from recordings of a linear response in MT in human (Rees et al. 2000; Händel et al. 2007) and monkey (Britten et al. 1993). Saturation of the contrast response function is thought to be the result of normalization, a canonical computation in cortex (Carandini and Heeger 2011; Baker and Wade 2017). If the response to motion coherence is linear, it might suggest that similar normalization does not apply. In fact, models of the V1 to MT circuitry include explicit normalization (Simoncelli and Heeger 1998) and the normalization strength alters whether the model predicts linear or saturating responses. This may account for the discrepancies of results; i.e. normalization may result in weak saturation of coherence response as we have found, in line with evidence from both humans (Rees et al. 2000; Costagli et al. 2014) and monkeys (Britten et al. 1993). In support of this idea is evidence that in the absence of a normal input from V1 the coherence response function in MT becomes more linear, possibly reflecting an increased input from subcortical regions whose coherence response is linear (Ajina et al. 2015). To clarify this we can again turn to behavior. Because the MT response has been linked to behavior (Katz et al. 2016) our model makes a testable prediction: under the assumption that the visual system performs signal detection subject to additive noise (Boynton et al. 1999) a saturating coherence function would predict worse discriminability of coherence at higher base levels of coherence.

To build out our quantitative framework we measured responses to stimuli of varying durations, down to those typically used in psychophysical experiments (e.g. 0.25 s) as well as at durations more typically used for bold> measurement (e.g. 4 s). Our results confirm many previous results showing that there exists a nonlinearity in the bold> response, such that shorter stimuli have a larger response than expected by temporal linearity (Boynton et al. 1996, 2012). Modeling our responses, we found that on average across cortical areas a doubling of the stimulus duration was associated with an increase in response of only 67% of the expectation of a linear model. Whether or not this is due to neural adaptation (Buxton et al. 2004) or saturation of the bold> signal (Friston et al. 1998) cannot be determined from our data. We also observed a slight difference in the duration effect across cortical areas. In V1 increasing duration results in a larger effect on the population response whereas MT showed a smaller than average response which could be a result of the more transient response in MT (Stigliani et al. 2017).

We also noted that any change in the stimulus, regardless of the type, amplitude, or duration resulted in what we refer to as a stimulus-onset response in all cortical areas. What is the nature of this response? Early recordings comparing bold> responses to electrophysiological recording suggest that the bold> signal may be thresholded at some minimum response even though neural activity continues to be modulated below that threshold (Logothetis et al. 2001). Another possibility is that the stimulus-onset response may be the result of a consistent trial structure causing anticipatory responses (Cardoso et al. 2012). In the latter case fitting a separate stimulus onset parameter to absorb this trial-structure related variance is appropriate to correctly estimate the population response from the bold> signal.

Our approach to making a parametric model of cortical response to motion visibility contrasts with more complex models, such as gabor-wavelet pyramids and deep convolutional networks (Kay et al. 2008; Yamins et al. 2014; Kay and Yeatman 2017) that are typically image-computable and thus can make detailed predictions of cortical response properties directly from images. A complete image computable model would implicitly contain our parametric model within it and seemingly obviate the need to parameterize stimulus visibility and its relationship to cortical response. Building such complex models is a worthy goal, however, we would note that much success in understanding visual cortex function has come from experiments which parametrically altered visual features, in particular features related to visibility. Consider the result of stimulus combination. When two gratings with different luminance contrast are presented the evoked response is not well captured by simple rules such as linear summation or winner-take-all (Busse et al. 2009) Instead, across a large range of parameter combinations the evoked response is well explained by normalization (Carandini and Heeger 2011). The canonical rule that an evoked response should be scaled by the total response of a region of cortex is easily understood in a parametric model, but far less intuitive in a complex one. Another low-dimensional parametric model is the population receptive field (Dumoulin and Wandell 2008; Wandell and Winawer 2015), which has been widely used to map and interpret the properties of retinotopic visual cortex, largely because of its simplicity. In general, low-dimensional quantitative frameworks like the one we have built can parameterize cortical response to key stimulus properties and by doing so, serve to make testable predictions for perceptual function. For example, our framework suggests that small variation in sensitivity across cortical areas might be used to separately determine the visibility of motion for different parameters. That is, a read-out of the visual representations could take advantage of the differences in feature sensitivity by differentially weighting V1 and MT for contrast discrimination and vice-versa for coherence discrimination.

Each parameter of motion visibility which we studied has been separately used to uncover the neural basis of different aspects of perception and perceptual decision making. The quantitative framework that we have proposed here shows that despite their similar effects on perception, contrast, coherence, and duration have distinct cortical representation at the level of populations. In studies of perception the effects of these parameters on cortical response should not be considered to be interchangeable. With our reference framework though one can now make changes in one parameter or the other and predict how this will affect human cortical response. In this way our predictive model is a key tool in furthering the goal of linking cortical response to perceptual behavior.

## Acknowledgments

We acknowledge the generous support of Research to Prevent Blindness and Lions Clubs International Foundation, and the Hellman Fellows Fund to J.L.G. This work was supported by the Stanford Center for Cognitive and Neurobiological Imaging. We particularly thank Steeve Laquitaine for assistance with data collection and Ian Eisenberg for continuous discussions and feedback. We thank Bob Dougherty for assistance with the high-resolution pulse sequence, Lynda Ichinaga for administrative support, and Guillaume Riesen, Akshay Jagadeesh, Minyoung Lee, and Kara Weisman for feedback on previous versions of this manuscript. We thank Bill Newsome for guidance and support throughout this project.

